# Scene memorability reflects representational distinctiveness within visual categories

**DOI:** 10.64898/2026.03.20.713124

**Authors:** Charlotte Atzert, Filip Děchtěrenko, Jiří Lukavský, Niko A. Busch

## Abstract

Some images are consistently remembered better than others, suggesting that memorability reflects intrinsic image properties. We tested whether within-category distinctiveness underlies this effect. Across three experiments (*N* = 477), participants categorized indoor scenes previously rated for subjective typicality and then completed recognition memory tests. Typical scenes were categorized faster and more accurately, but were remembered worse and showed a more liberal response bias than atypical scenes. These opposing effects were robust across categories. To link subjective typicality to visual representations, we quantified image distinctiveness using a convolutional neural network (CNN). Across layers, CNN-derived distinctiveness closely tracked human typicality judgments and predicted both categorization speed and memorability, with strongest effects in higher, semantic layers. Critically, the memory advantage for atypical scenes persisted even when most images were atypical, ruling out rarity within the experimental context. Together, the results show that intrinsic scene memorability reflects an image’s position within a category-specific representational space.

## Introduction

One of the central aims of memory research is to understand why some events are remembered while others are forgotten. Although personal and situational factors such as attention, motivation, and emotional engagement strongly influence memory, the content of an event also contributes substantially to memorability. In the domain of long-term memory for scene images, a growing body of work shows that some scenes are consistently remembered or forgotten across observers [1, 2], implying that memory performance is partly determined by properties intrinsic to the image rather than by individual experience or task context. This intrinsic memorability accounts for a substantial proportion of variance in performance [3], suggesting that it reflects fundamental mechanisms of human memory and raising the question of which image properties give rise to this consistency.

What intrinsic image features drive memorability? Low-level features such as luminance, color, saturation, or the number of objects are at best weak predictors of memorability, and any observed correlations may largely reflect their association with higher-level properties [4]. For example, the modest advantage of images with a warm hue may arise from the higher memorability of images containing faces. Several mid-to high-level properties – including the presence of a salient fore-ground object, moderate clutter, scenes with people, close-up faces, animals [4], and good perceptual organization [5] – have also been shown to increase memorability. In addition, memorability varies systematically across scene categories, with urban and indoor scenes remembered better than landscapes [6]. Importantly, however, memorability also varies reliably among exemplars within the same category [6, 7]. Although memorability cannot be explained by any single image attribute, it can be predicted from combinations of features extracted by deep neural networks [8]. For example, the ResMem model [9] predicts human memorability with high accuracy. However, while such models demonstrate a robust statistical relationship between image features and memorability, the psychological mechanisms underlying these predictions remain poorly understood. How do humans process and encode images in ways that give rise to these regularities?

We propose that an important factor influencing intrinsic image memorability is an image’s position within the representational structure of a stimulus set or a category, that is, its typicality or distinctiveness. These terms describe opposing ends of a continuum: typical items cluster near the prototype, whereas distinctive items are relatively distant from it. Distinctiveness thus broadly refers to the degree to which an item’s perceptual or conceptual features deviate from those of other items or category members [10, 11]. Classic work by von Restorff [12] showed that distinctive items that stand out from their context (e.g., a red letter among black letters) are remembered better than common items. Subsequent research distinguished between primary distinctiveness, defined relative to a specific set of items in an experiment, and secondary distinctiveness, referring to items that are atypical or “peripheral members of a natural category” [13, p. 530].

Consistent with the view that semantic categories are continuous rather than discrete [14], exemplars vary in the degree to which they are judged typical of their category. Numerous studies have shown that typicality – or conversely, secondary distinctiveness, referring to items that are atypical members of a category – influences recognition memory (with less consistent effects on recall; [15]). For example, atypical or “out-of-place” objects (e.g., a mailbox in a kitchen) are remembered better than typical objects [16], and faces judged as atypical are recognized more accurately than typical faces [17–19].

A useful way to formalize typicality and distinctiveness is to conceptualize semantic categories as multidimensional representational spaces, where each dimension represents a feature relevant for distinguishing between exemplars or categories, and items are represented as points or vectors. In such spaces, typical exemplars cluster near the center, whereas distinctive exemplars are more distant and isolated [15, 20, 21]. This geometric interpretation underlies influential theories of categorization, which represent exemplars and prototypes as vectors in multi-feature spaces [22]. Within this framework, response times in category verification tasks are thought to reflect the distance between an exemplar and a category prototype, that is, the degree of feature overlap [23]. Accordingly, typical exemplars are categorized more rapidly and accurately than atypical exemplars [14, 24, 25].

Recent studies have extended this representational-space approach to natural scenes by quantifying distinctiveness using feature representations derived from deep convolutional neural networks (CNNs). However, results remain mixed: some studies report higher memorability for scenes with distinctive CNN features [6, 21], whereas others find better memory for scenes with more common features [26, 27].

Against this background of mixed empirical findings, the present study aimed to clarify the role of scene typicality – particularly within-category differences in typicality – in determining intrinsic scene memorability. To isolate effects of exemplar typicality, we tested memory for a large set of exemplars from three indoor scene categories, thereby controlling for differences in memorability between categories [27] that can confound studies including multiple categories. Exemplar typicality was assessed using both subjective typicality ratings and analyses of image features derived from a CNN. In addition, participants completed a speeded category verification task in which they judged whether a scene belonged to a specific category.

Assuming that the representational space underlying scene categorization overlaps with that supporting scene memory, we predicted that typical exemplars would be categorized more efficiently but remembered less accurately than atypical exemplars. By testing these predictions, the present study aims to elucidate the cognitive mechanisms underlying intrinsic image memorability and its computational basis, linking scene representations in artificial neural networks to human perceptual and conceptual representations.

## Experiment 1

### Methods

#### Participants

Data were collected from 92 participants (30 in the lab, 62 online; 65 female; age range 18–66 years, *M* = 27; 79 right-handed). Two participants were excluded for having fewer than 10 valid trials per typicality condition in the categorization task (from a total of 126 target trials), and 20 were excluded for making false alarms to more than one catch trial in the memory task, leaving data from 70 participants for analysis. All participants provided written informed consent and received course credit or monetary compensation. The study was approved by the Ethics Committee of the Faculty of Psychology and Sports Sciences, University of Münster (approval #2024-13-NB).

**Figure 1:**
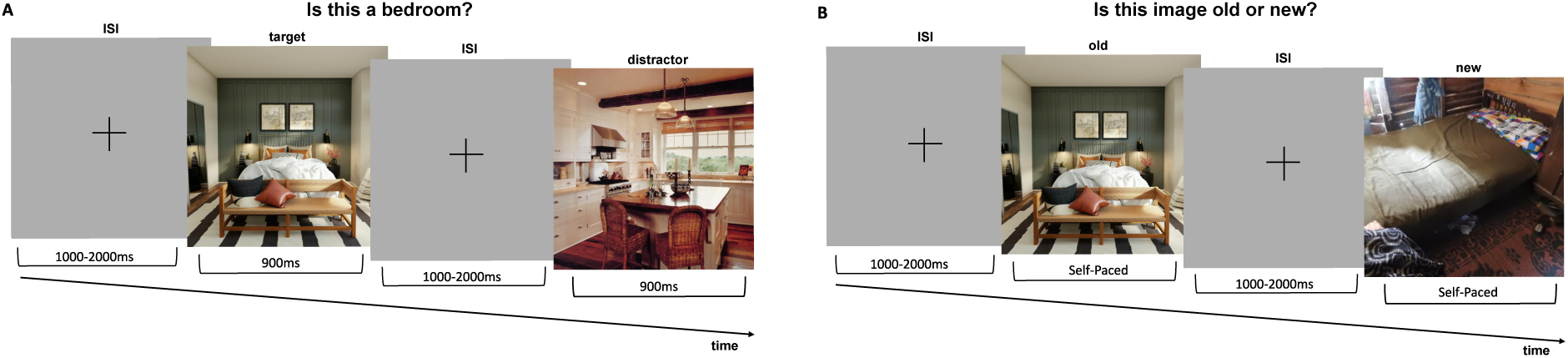
Illustration of a trial sequence. (A) Categorization task: Each trial began with a fixation cross (1000–2000 ms), followed by an image presented for 900 ms. Participants indicated as quickly as possible whether the image belonged to the target category. (B) Memory task: Participants indicated whether the image was old or new; responses were self-paced.

#### Stimuli and Apparatus

The experiment was implemented in PsychoPy version 2024.1.5 [28] and run locally in the lab (Python code) and online via Pavlovia.org (JavaScript code).

Stimuli were selected from a pool consisting of 1,518 images from three indoor scene categories (499 bedrooms, 727 kitchens, and 292 living rooms). Images were selected from public sources (https://unsplash.com, https://www.pexels.com, https://pixabay.com), or established scene databases (change blindness database, [29]; SUN database, [30]). They were preselected to span a wide range of typicality regarding the objects present, scene layout, framing, and perspective. All images were free of people, imaging artifacts, and text annotations. An independent sample of 310 participants rated all scenes for semantic typicality (“How well does the label x describe this image?”) and perceptual typicality (“How many images of x look similar to this one?”) on a 10-point scale. Since the two typicality measures were highly correlated, we quantified typicality in the present study as the mean of both measures (Figure 3, top row). Stimuli and typicality ratings are available at https://zenodo.org/records/18935615.

#### Procedure

##### Categorization task

Each categorization block began with the presentation of a text label indicating the target scene category for that block (kitchen, bedroom, or living room), followed by a sequence of scene images. Each trial consisted of a fixation cross presented for 1,000–2,000 ms (randomized per trial) and a scene image shown for 900 ms. Participants responded as quickly as possible using the “f” and “j” keys to indicate whether the image was an exemplar of the target category or a distractor. A key response was required to proceed to the next trial.

Each block contained 21 target images and 7 distractors drawn from the remaining two categories. Target images were randomly sampled from the stimulus pool, with the restriction that seven had typicality ratings below the median and fourteen above it. Distractors were sampled randomly regardless of typicality. Each category served as the target in two blocks (six blocks in total), and block order was counterbalanced across participants. All experimental blocks were preceded by a practice block with the same structure, using images from two unrelated categories (airports and train stations).

##### Memory task

Following the six categorization blocks, all 126 target images were presented again together with new foil images from the same categories that had not been shown before. Each trial began with a fixation cross shown for 1,000–2,000 ms (randomized per trial), followed by a scene image that remained on screen until a response was made. For each image, participants indicated whether it was old or new and rated their confidence (“sure” or “maybe”). Participants were not instructed to respond quickly; images remained on screen until a response was made.

The proportion of foils was .33. Distractor images from the categorization task were not repeated in the memory task. To keep the lag between image occurrences approximately constant across all items, images were presented blockwise, using the same block order as in the categorization task. Within each block, the presentation order was randomized. Additionally, five catch trials featuring novel scenes from unrelated categories (bathroom, office, study, and nursery) were randomly interspersed within the memory blocks.

#### Analysis

Response time (RT) analysis in the categorization task was restricted to correct responses. RTs faster than 200 ms or slower than 3000 ms were excluded.

Accuracy in both tasks was quantified using signal detection measures of sensitivity (*d*^′^) and response criterion (*c*) [31]. To avoid extreme hit or false-alarm rates of 1 and 0, 0.5 was added to both the number of hits and false alarms, and 1 to both the number of old and new items before computing hit and false-alarm rates [32].

Response times and accuracy were analyzed in separate 2 x 3 repeated measures analyses of variance (ANOVA) as implemented in the afex package (version 1.3; [33]) for R [34] with factors *typicality* (high, low) and *category* (bedroom, kitchen, living room). Degrees of freedom and p-values were Greenhouse–Geisser corrected where appropriate. Pairwise post-hoc contrasts were computed using the emmeans package (version 1.6.3; [35]).

## Results

Categorization and recognition memory performance is summarized in Figure 2A.

**Figure 2:**
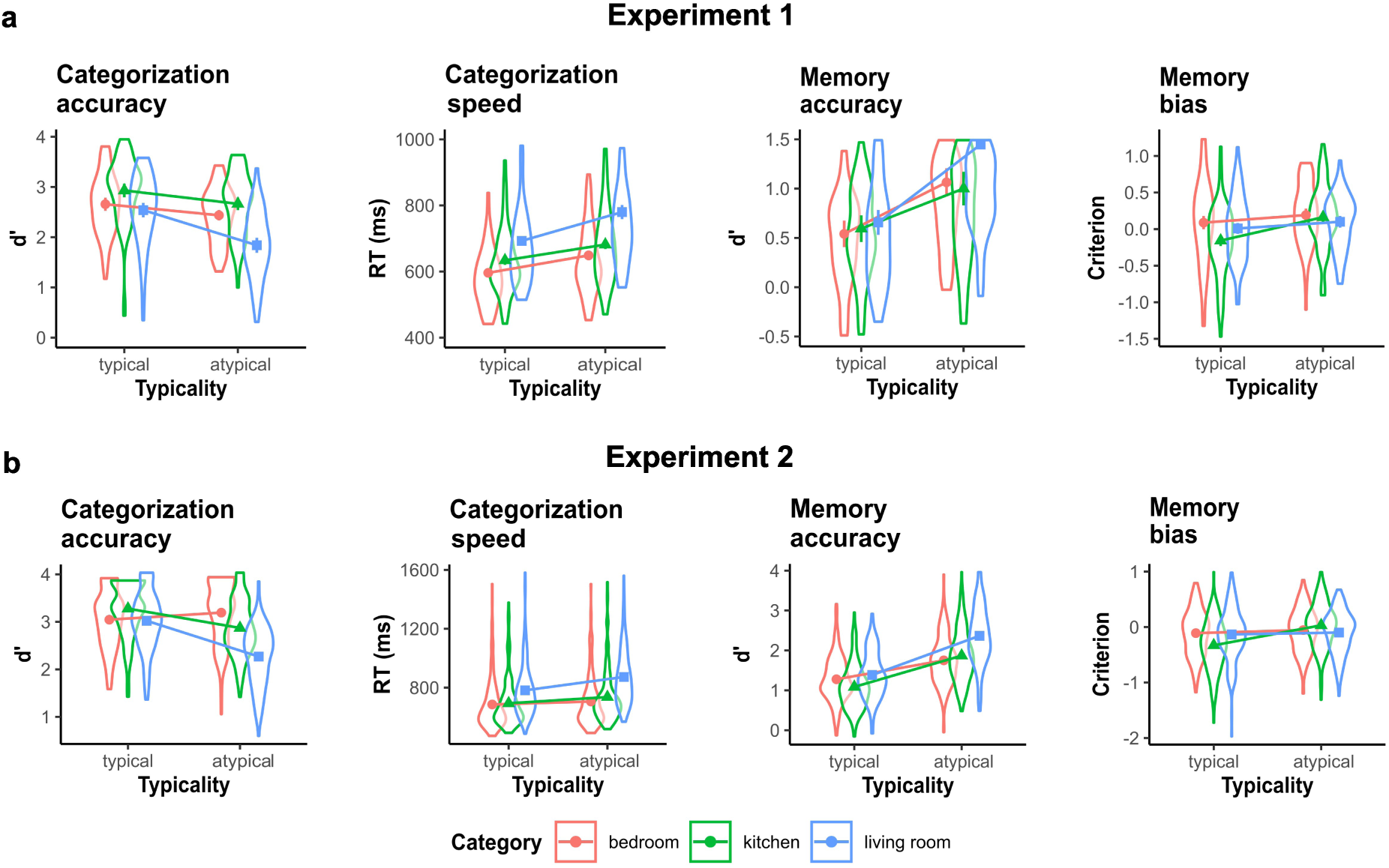
Categorization and recognition memory performance in Experiments 1 and 2. (A) Experiment 1: Categorization was more accurate and faster for typical than for atypical scenes. In contrast, recognition memory was less accurate for typical scenes. The recognition criterion was lower for typical scenes, indicating a more liberal bias to judge typical images as “old.” (B) Experiment 2: Overall memory accuracy was higher than in Experiment 1, but the same typicality effects on categorization and recognition memory were observed.

**Figure 3:**
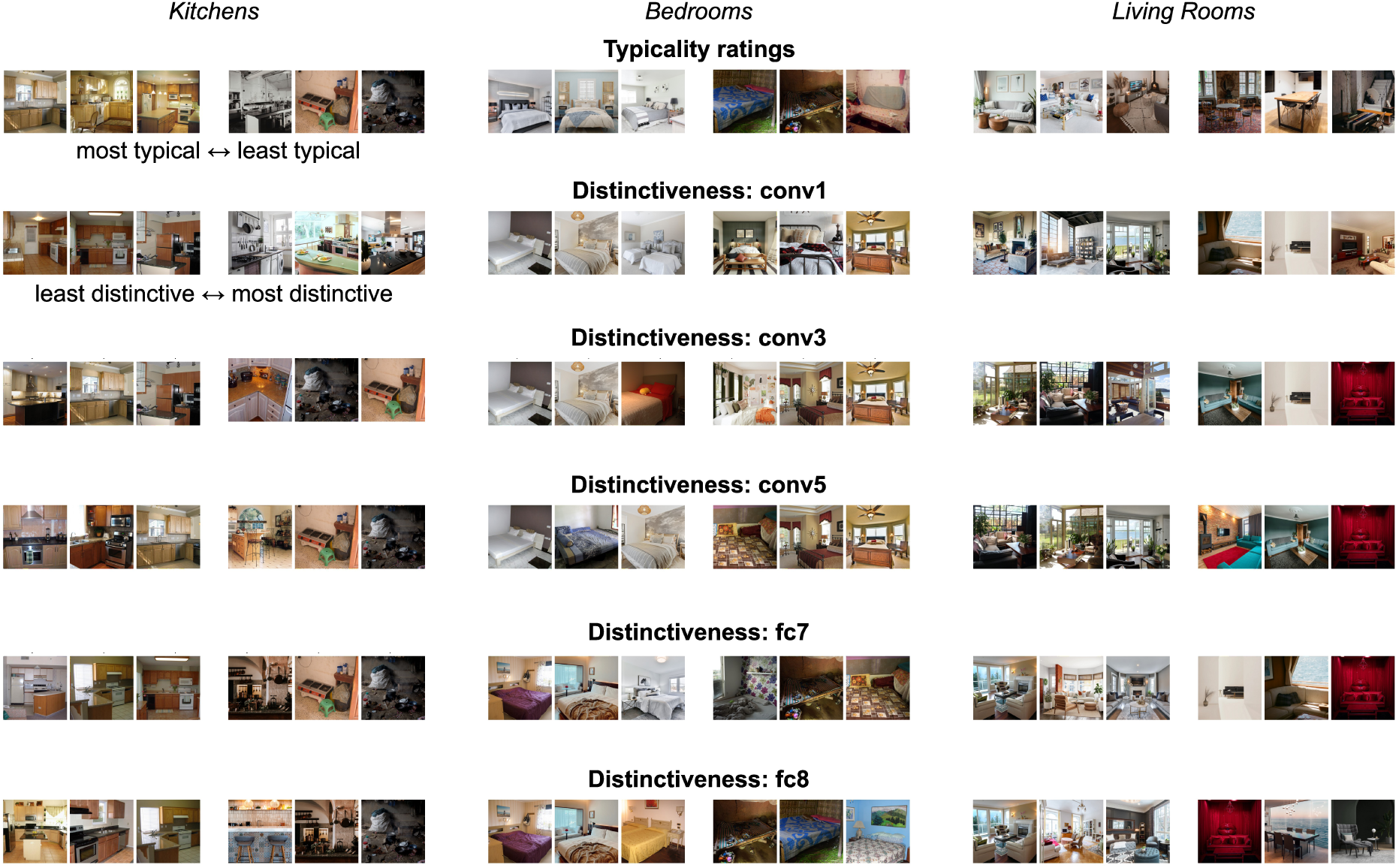
Examples of scene exemplars, sorted by human typicality ratings and CNN-based feature distinc-tiveness, respectively. Top row shows scenes with the highest and lowest human typicality ratings (left: most typical; right: least typical). Rows below show scenes with the lowest and highest feature distinctiveness (left: least distinctive; right: most distinctive), computed separately for different layers of the deep convolu-tional neural network (conv1, conv3, conv5, fc7, fc8). Distinctiveness was calculated within scene category, such that higher values indicate greater dissimilarity from other exemplars of the same category. A general correspondence between human typicality judgments and CNN-based distinctiveness can be observed, although distinctiveness showed a clear progression across layers in how it aligns with typicality judgments. Distinctiveness derived from early convolutional layers (conv1-conv3), which primarily capture low-level visual features such as edges, textures, and local contrast, produced relatively subtle differences between typical and atypical exemplars. In contrast, distinctiveness computed from higher convolutional and fully connected layers (conv5, fc7, fc8) yielded increasingly pronounced and semantically coherent distinctions, with atypical scenes appearing more visually and conceptually idiosyncratic.

### Categorization Accuracy

Categorization was more accurate for typical targets than for atypical targets (*F* (1, 69) = 202.03, *p* < .001, 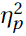 = .75). Moreover, accuracy was generally highest for kitchen scenes, intermediate for bedroom scenes, and lowest for living room scenes (*F* (1.79, 123.18) = 27.32, *p* < .001, 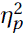 = .28). A significant interaction (*F* (2, 138) = 12.13, *p* < .001, 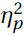 = .15) indicated that the typicality effect on accuracy was strongest for living room scenes (*t* (69) = 11.71, *p <* .001), intermediate for kitchen scenes (*t* (69) = 8.59, *p <* .001), and lowest for bedroom scenes (*p*(69) = 6.30, *p* < .001).

### Categorization Speed

Categorization was faster for typical targets than for atypical targets (*F* (1, 69) = 156.69, *p* < .001, 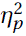 = .69). Moreover, responses were generally faster for kitchen scenes, intermediate for bedroom scenes, and slowest for living room scenes (*F* (1.82, 125.76) = 79.40, *p* < .001, 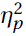 = .54). A significant interaction (*F* (1.77, 121.89) = 7.16, *p* = .002, 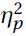 = .09) indicated that the typicality effect on categorization speed was strongest for living room scenes (*t* (69) = −8.90, *p <* .001), intermediate for bedroom scenes (*t* (69) = −7.34, *p <* .001), and lowest for kitchen scenes (*t* (69) = −6.41, *p <* .001).

### Recognition Accuracy

Recognition was more accurate for atypical targets than for typical targets (*F* (1, 69) = 96.49, *p* < .001, 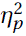 = .58). Moreover, recognition was generally most accurate for living room scenes, intermediate for bedroom scenes, and lowest for kitchen scenes (*F* (2, 138) = 7.68, *p* < .001, 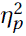 = .10). A significant interaction (*F* (2, 138) = 4.25, *p* = .016, 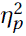 = .06) indicated that the typicality effect on accuracy was strongest for living room scenes (*t* (69) = −9.19, *p <* .001), intermediate for bedroom scenes (*t* (69) = −5.74, *p <* .001), and lowest for kitchen scenes (*t* (69) = −3.60, *p <* .001).

### Recognition Criterion

The recognition criterion was negative for typical scenes (*M* = –0.02) and positive for atypical scenes (*M* = 0.15), indicating that participants were biased to judge typical scenes as “old.” This liberal bias for typical scenes was confirmed by a main effect of typicality, (*F* (1, 69) = 22.28, *p* < .001, 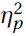 = .24). Criterion also differed across scene categories, being most liberal for kitchen scenes, intermediate for living rooms, and most conservative for bedrooms (*F* (2, 138) = 5.62, *p* = .005, 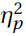 = .08). A significant interaction between typicality and category (*F* (1.81, 125.07) = 5.65, *p* = .006, 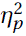 = .08) showed that the typicality effect on criterion was strongest for kitchen scenes (*t* (69) = −6.09, *p <* .001), but not significant for living room scenes (*t* (69) = −1.86, *p <* .067), or bedroom scenes (*t* (69) = −1.53, *p <* .131).

## Discussion

In Experiment 1, we examined how scene typicality influences categorization and recognition memory. Typical scenes were categorized more quickly and accurately than atypical scenes, whereas recognition memory showed the opposite pattern: atypical scenes were remembered better than typical ones. Participants also adopted a more liberal response criterion for typical scenes, indicating a greater tendency to falsely recognize new typical images as previously seen, consistent with a stronger sense of familiarity. Importantly, these opposing effects on categorization and memory argue against a low-level explanation such as image quality, which would be expected to impair both tasks in the same direction. Although the magnitude of the typicality effects varied across scene categories, the pattern was consistent in all three, suggesting that atypical scenes are processed as more distinct category members.

The categorization results align with previous findings showing slower and less accurate categorization for atypical exemplars [14, 24, 25]. Such effects have been interpreted within different theoretical frameworks of category representation and decision-making. Prototype theories propose that observers abstract a central tendency from category exemplars and categorize new items based on their similarity to this prototype [36]. Other accounts emphasize similarity to the distribution of category features [37] or to stored exemplars [38]. Despite their differences, these models converge on the prediction that categorization becomes slower as the representational distance between a target and the bulk of category members increases. Accordingly, the strong relationship between subjective typicality ratings and categorization speed supports the interpretation of these ratings as indexing representational distinctiveness.

The memory results showed the complementary pattern: atypical scenes were remembered better than typical ones. This advantage for atypical exemplars parallels prior findings in other domains, including words [39] and faces [17–19]. Together with their slower categorization, this pattern suggests that atypical scenes occupy more isolated positions in representational space, making them less confusable with other stored items and therefore easier to recognize [21]. The combination of faster categorization but poorer recognition for typical exemplars, and the reverse for atypical ones, closely mirrors classic findings on face typicality [24].

In sum, the complementary effects of typicality on categorization and recognition memory support our hypothesis that intrinsic scene memorability is, at least in part, determined by a scene’s typicality and its location in representational space relative to other exemplars of the same category. Nonetheless, several aspects of the design of Experiment 1 limit the strength of this conclusion and were explicitly addressed in Experiment 2.

### Experiment 2

Experiment 1 showed that atypical scenes were categorized more slowly but recognized more accurately than typical ones. However, these group-level effects do not establish whether an individual image’s typicality jointly predicts its categorization speed and memorability. Addressing this question requires image-by-image analyses that correlate categorization and recognition performance across individual scenes. Moreover, directly testing the hypothesis that atypical scenes are more distinctive in representational space requires quantifying distinctiveness for each image based on its visual features and relating this measure to behavioral performance. Such analyses demand a consistent number of observations per image. Because Experiment 1 sampled stimuli randomly from a larger pool, resulting in few and uneven observations per image, Experiment 2 presented the same set of scenes to all participants, using a more balanced selection of typical and atypical exemplars.

In addition, overall recognition performance in Experiment 1 was low, with several participants performing at or near chance. In Experiment 2, each categorization block was immediately followed with a recognition test for the images from that block only, thereby shortening the encoding–retrieval delay and reducing interference between items [11].

To quantify image-level distinctiveness, we extracted visual features from a deep convolutional neural network. CNNs are hierarchical models that transform pixel values into increasingly abstract feature representations, capturing low-level features such as edges and textures in early layers and higher-level, semantically meaningful information in later layers [40]. These representations show important parallels to the primate visual system, with early layers resembling feature detectors in early visual cortex and later layers corresponding to higher-level visual areas [41, 42]. Consistent with this view, activity patterns from higher CNN layers predict neural responses in monkey inferotemporal cortex and human ventral visual cortex [43, 44]. Although CNNs are not a complete model of human vision [45], feature similarities derived from CNNs correspond well with human perceptual similarity judgments [46, 47], making them a useful tool for estimating scene distinctiveness.

Koch *et al.* [26] found that distinctiveness in early CNN layers predicted higher memorability, whereas distinctiveness in later, category-related layers predicted lower memorability. This counter-intuitive pattern challenges the assumption that atypical images should be more memorable and raises important questions about how distinctiveness at different representational levels contributes to image memorability. Accordingly, Experiment 2 examined how CNN-derived distinctiveness across multiple layers relates to human typicality judgments and how both measures predict memorability across a wide range of exemplars within the same semantic category.

#### Methods

Methods were identical to experiment 1 except for the following changes.

#### Participants

We collected data from 211 participants (139 female; age range 19-77, mean 27 years; 162 right-handed). Data from 6 participants were excluded because they included less than 10 correct trials per typicality condition within the range from 200 to 2000 ms. Thirty-five participants were excluded because they made false alarms to more than 1 catch trial. Overall, the analysis included data from 174 participants.

#### Stimuli and Apparatus

The experiment was only conducted online. The same selection of scene images was used for all participants, but with variable assignments to the experimental conditions. For each scene category, the images in the image pool were ranked by typicality ratings and binned into deciles. For half of the participants, the target images in the categorization task were selected for each block as the image with the lowest and highest typicality values within each decile (20 targets in total). Twenty additional distractor images from non-target categories were selected whose typicality ratings were closest to the mean typicality of their respective categories. Foil items for the memory task were selected as the two images with typicality ratings closest to the mean typicality of each bin (20 foils in total). For the other half of the participants, the targets and foils were reversed so that each image was used as an old target and as a new foil in half of the participants.

#### CNN features and distinctiveness metrics

To test the hypothesis that scene typicality and memorability are linked to the unusualness of a scene’s visual features, we analyzed all images using layer-wise feature representations derived from a deep convolutional neural network. Specifically, we used a pretrained AlexNet model [40], implemented in PyTorch [48] and trained on the Places365 dataset, which contains 1.8 million high-resolution natural images spanning 365 scene categories [49]. The network comprises eight sequential layers: five convolutional layers followed by three fully connected layers.

As an initial validation that the network encoded the scene categories (kitchen, bedroom, living room) in a manner comparable to human observers, we correlated human typicality ratings with the probability that the network classified each image as its correct category, based on the final classification layer (fc8). Across all images, this correlation was r = 0.517, p < .001. Although the absolute magnitude of this effect should be interpreted cautiously – given differences between the network’s fine-grained category structure (e.g., bedroom vs. bedchamber) and the broader category labels likely used by human observers – the result indicates that scenes judged as more typical by humans were also more likely to be correctly categorized by the network [50]. We therefore asked whether this correspondence could be explained by the distinctiveness of image representations.

Because CNN feature representations become increasingly complex and category-specific across layers [41, 42], we examined the emergence of image distinctiveness across the network hierarchy. Analyses focused on convolutional layers 1, 3, and 5, the penultimate fully connected layer (fc7), and the final classification layer (fc8). For convolutional layers, the feature maps were flattened into a one-dimensional feature vector.

Image distinctiveness was quantified separately for each layer by computing pairwise Pearson correlations between feature vectors of all image pairs within the same scene category, yielding category-specific image-by-image similarity matrices. For each image, mean representational similarity was defined as the average of its Fisher z–transformed correlations with all other images from the same category. Distinctiveness was then computed as the inverse of this mean similarity, such that higher values indicate greater representational distance from other category members (Figure 3). This measure corresponds to the DCNN-based typicality metric in Kramer *et al.* [27] and to the inverse of the image similarity metric proposed by Koch *et al.* [26]. Finally, we correlated layer-wise distinctiveness to human typicality ratings, categorization response times, and memory performance (Figure 5).

**Figure 4:**
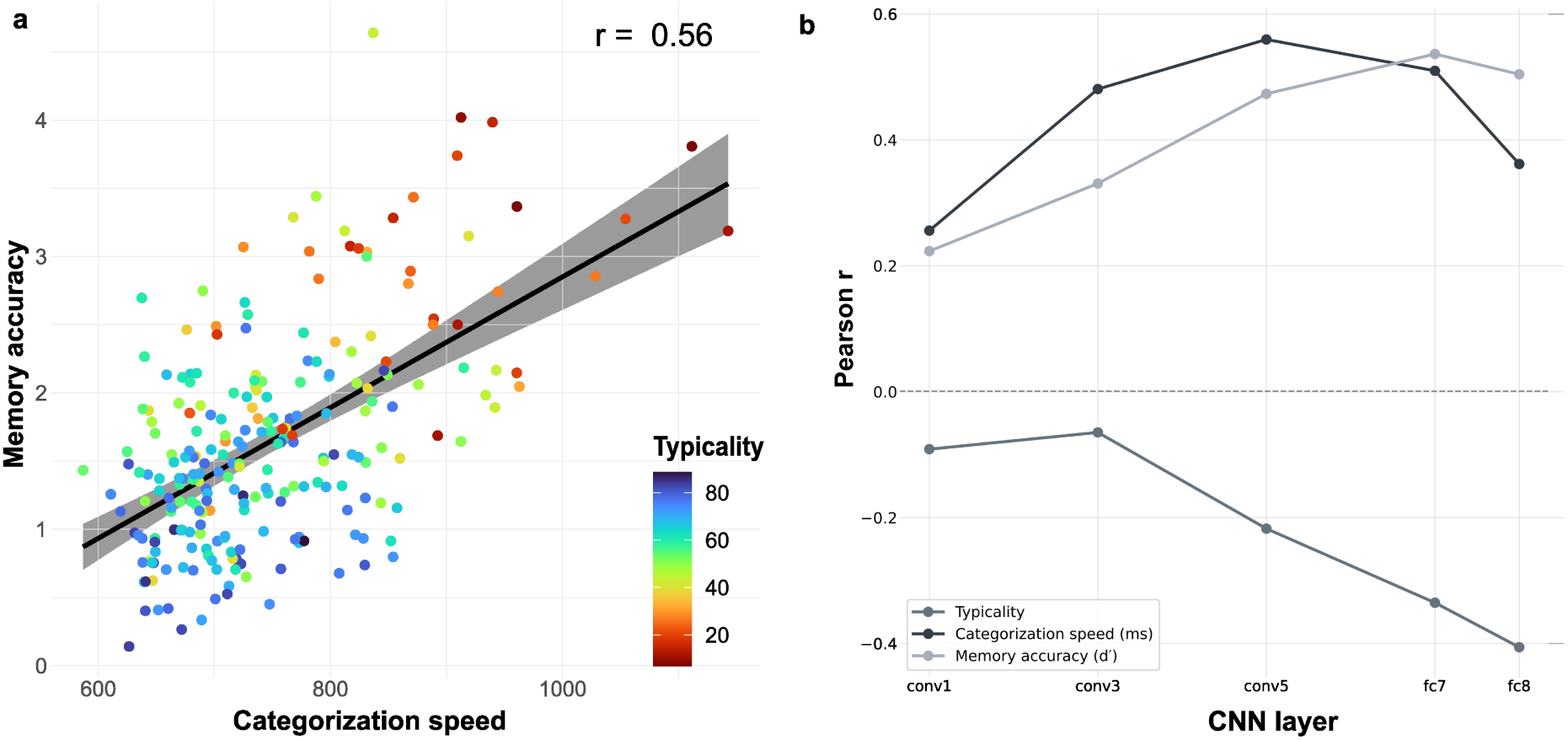
(A) In Experiment 2, categorization speed predicted recognition accuracy, and this relationship was mediated by scene typicality. (B) Overview of correlations between behavioral measures and CNN-derived distinctiveness. Categorization speed showed the strongest association with features from convolutional layer 5, with weaker correlations in earlier and later layers. In contrast, typicality and recognition accuracy exhibited a more consistent increase in correlation strength across layers, peaking in the later layers.

**Figure 5:**
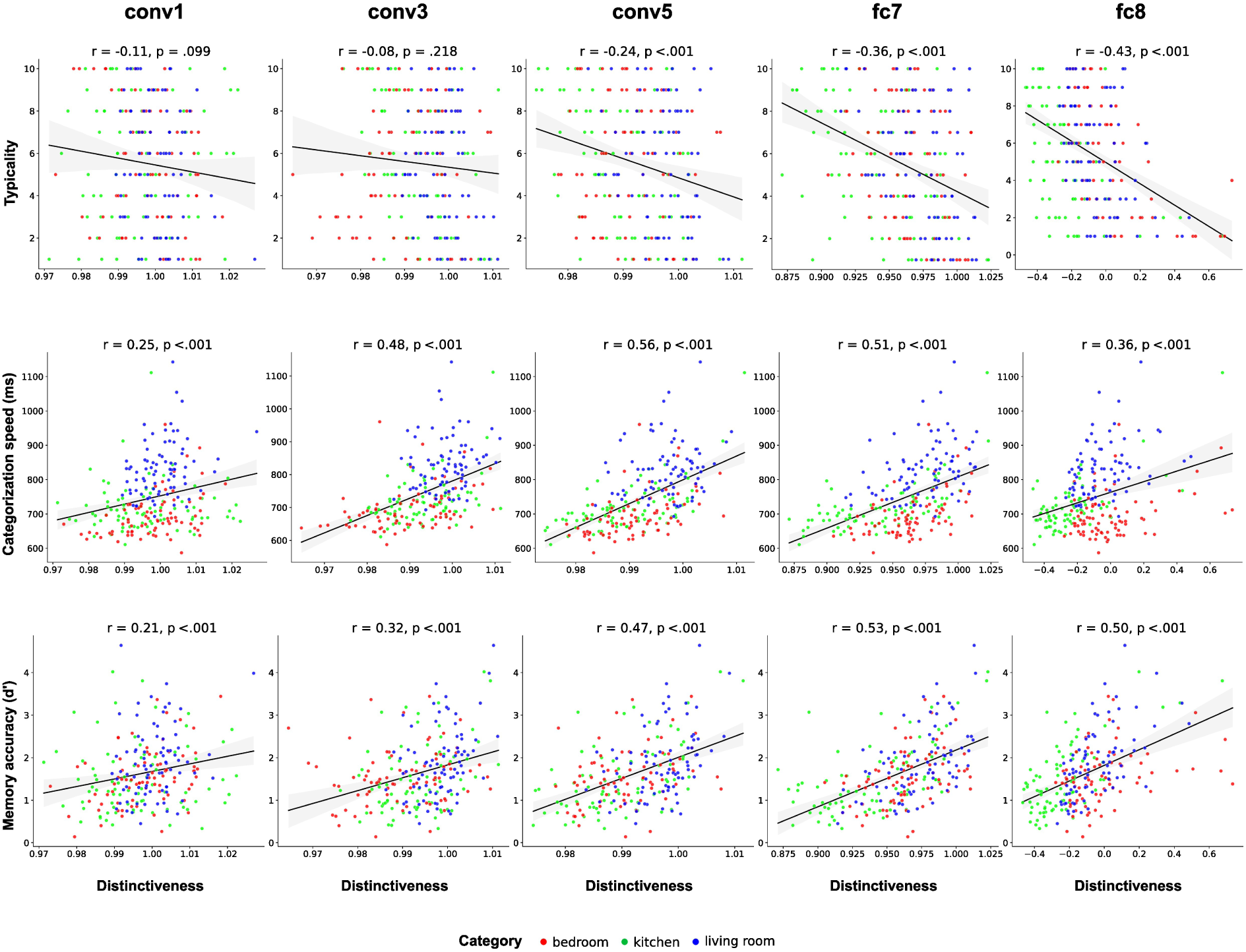
Correlations between CNN-based distinctiveness and behavioral measures. From left to right, distinctiveness was computed from features in convolutional layers conv1, conv3, and conv5, the fully connected layer fc7, and the final classification layer fc8. Top row: More distinctive images were judged as less typical, with correlation strength increasing across layers. Middle row: More distinctive images were categorized more slowly, with the strongest relationship observed for features from conv5. Bottom row: More distinctive images were remembered more accurately, with correlations increasing across layers.

Code for extracting and applying deep neural network features was generated with the assistance of a LLM, subsequently verified and adapted.

#### Procedure

Categorization blocks included 20 targets and 20 distractors each. Note that the categorization task in experiment 1 contained substantially more targets than distractors, possibly biasing participants to respond “target” even for ambiguous images. Each categorization block was followed immediately by a memory block featuring all 20 target images from the preceding categorization block, 20 new foils of the target category, and one catch trial from a category that was not used as a target nor distractor in any block (stairwells or supermarkets).

## Results

Experiment 2 replicated all main findings from experiment 1 (Figure 2B). Specifically, typical scenes were categorized faster and more accurately than typical images. While overall memory performance was higher than in experiment 1, recognition was again more accurate and bias was more liberal for typical than for atypical images (see table 1 and 2 for details).

**Table 1:**
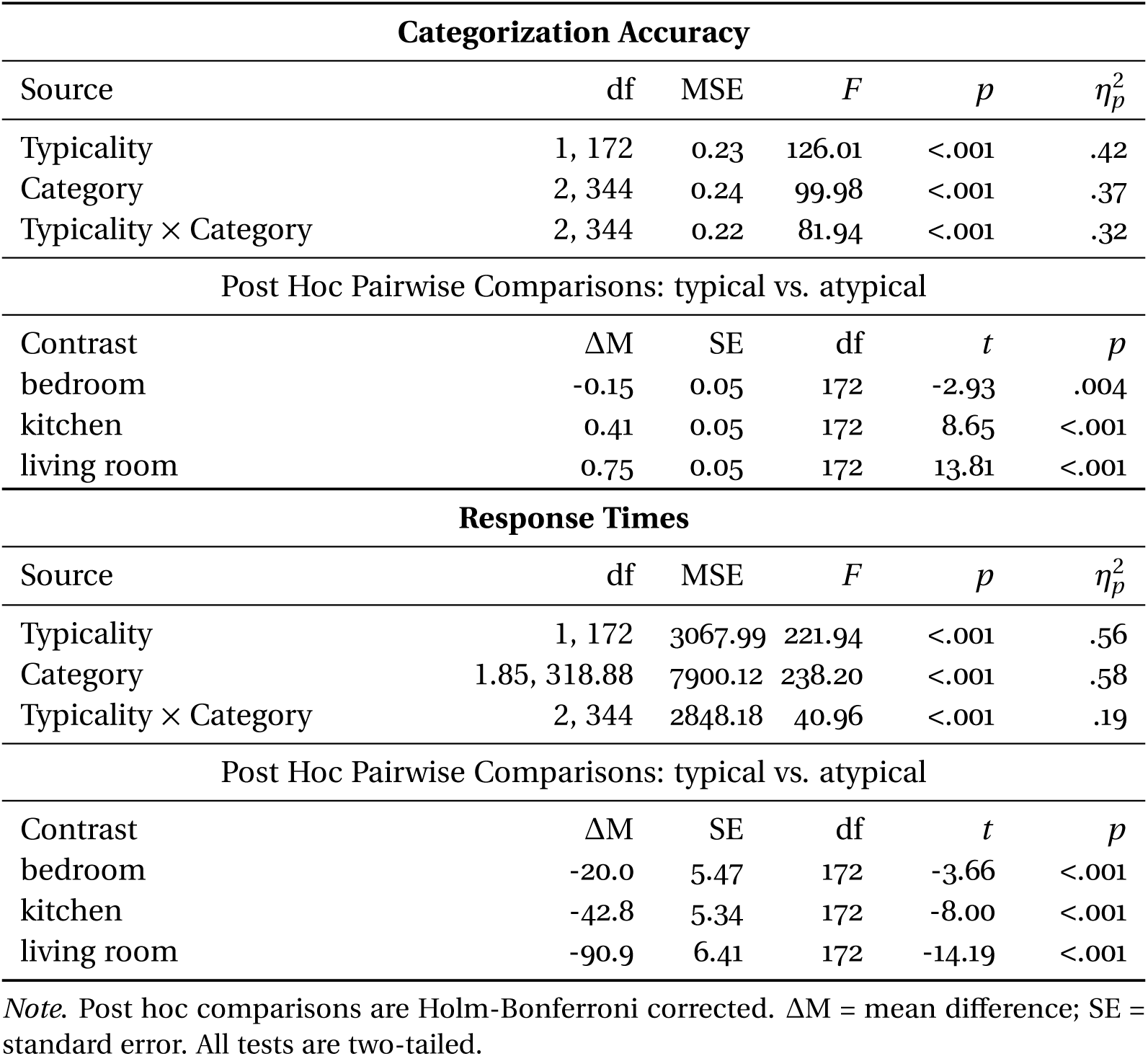
Two-Way ANOVA Results for the Categorization Task (Experiment 2)

| **Categorization Accuracy** |  |  |  |  |  |
| --- | --- | --- | --- | --- | --- |
| Source | df | MSE | *F* | *p* | $\eta_p^2$ |
| Typicality | 1, 172 | 0.23 | 126.01 | <.001 | .42 |
| Category | 2, 344 | 0.24 | 99.98 | <.001 | .37 |
| Typicality × Category | 2, 344 | 0.22 | 81.94 | <.001 | .32 |
| Post Hoc Pairwise Comparisons: typical vs. atypical |  |  |  |  |  |
| Contrast | $\Delta M$ | SE | df | *t* | *p* |
| bedroom | -0.15 | 0.05 | 172 | -2.93 | .004 |
| kitchen | 0.41 | 0.05 | 172 | 8.65 | <.001 |
| living room | 0.75 | 0.05 | 172 | 13.81 | <.001 |
| **Response Times** |  |  |  |  |  |
| Source | df | MSE | *F* | *p* | $\eta_p^2$ |
| Typicality | 1, 172 | 3067.99 | 221.94 | <.001 | .56 |
| Category | 1.85, 318.88 | 7900.12 | 238.20 | <.001 | .58 |
| Typicality × Category | 2, 344 | 2848.18 | 40.96 | <.001 | .19 |
| Post Hoc Pairwise Comparisons: typical vs. atypical |  |  |  |  |  |
| Contrast | $\Delta M$ | SE | df | *t* | *p* |
| bedroom | -20.0 | 5.47 | 172 | -3.66 | <.001 |
| kitchen | -42.8 | 5.34 | 172 | -8.00 | <.001 |
| living room | -90.9 | 6.41 | 172 | -14.19 | <.001 |

*Note.* Post hoc comparisons are Holm-Bonferroni corrected. $\Delta M$ = mean difference; SE = standard error. All tests are two-tailed.

**Table 2:** Two-Way ANOVA Results for the Memory Task (Experiment 2)

| **Memory Accuracy** |  |  |  |  |  |
| --- | --- | --- | --- | --- | --- |
| Source | df | MSE | *F* | *p* | $\eta_p^2$ |
| Typicality | 1, 172 | 0.23 | 625.86 | <.001 | .78 |
| Category | 2, 344 | 0.24 | 70.79 | <.001 | .29 |
| Typicality × Category | 2, 344 | 0.21 | 28.09 | <.001 | .14 |
| Post Hoc Pairwise Comparisons: typical vs. atypical |  |  |  |  |  |
| Contrast | $\Delta M$ | SE | df | *t* | *p* |
| bedroom | -0.47 | 0.05 | 172 | -10.29 | <.001 |
| kitchen | -0.78 | 0.05 | 172 | -15.74 | <.001 |
| living room | -0.98 | 0.05 | 172 | -18.77 | <.001 |
| **Memory Criterion** |  |  |  |  |  |
| Source | df | MSE | *F* | *p* | $\eta_p^2$ |
| Typicality | 1, 172 | 0.09 | 64.24 | <.001 | .27 |
| Category | 2, 344 | 0.07 | 5.42 | .005 | .03 |
| Typicality × Category | 2, 344 | 0.05 | 52.77 | <.001 | .23 |
| Post Hoc Pairwise Comparisons: typical vs. atypical |  |  |  |  |  |
| Contrast | $\Delta M$ | SE | df | *t* | *p* |
| bedroom | -0.05 | 0.03 | 172 | -1.91 | .058 |
| kitchen | -0.36 | 0.03 | 172 | -13.80 | <.001 |
| living room | -0.03 | 0.03 | 172 | -1.20 | .230 |

*Note.* Post hoc comparisons are Holm-Bonferroni corrected. $\Delta M$ = mean difference; SE = standard error. All tests are two-tailed.

An image-by-image analysis revealed that images categorized more rapidly were remembered more accurately (*r* = 0.56, *p* < .001; Figure 4A). To relate computational estimates of scene distinctiveness to human behavior, we correlated distinctiveness at each CNN layer with the image-wise averages of typicality, categorization speed, and recognition accuracy. These correlations were significant at all layers and generally increased in strength from early to higher layers (Figure 4B).

Specifically, greater distinctiveness was associated with lower typicality ratings across layers. This negative correlation was weak in early convolutional layers but increased across the network hierarchy, reaching its strongest expression in the fully connected and classification layers (Figure 5, top row). Across layers, greater distinctiveness was associated with slower categorization. This relationship was strongest for distinctiveness derived from conv5 features, with weaker correlations in earlier convolutional layers and later fully connected layers (Figure 5, middle row). Across layers, greater distinctiveness was associated with higher recognition accuracy. The strength of this positive correlation increased progressively from early to higher layers (Figure 5, bottom row).

## Discussion

Experiment 2 replicated all central findings of Experiment 1. Typical scene exemplars were categorized more rapidly and more accurately, yet were remembered less accurately than atypical exemplars. Crucially, image-by-image analyses revealed that an image’s categorization speed predicted its memory accuracy, and that this relationship was mediated by typicality.

Experiment 2 further tested whether this behavioral dimension of typicality reflects an image’s position in representational space by quantifying image distinctiveness using deep neural network features. Across CNN layers, greater distinctiveness was consistently associated with lower typicality ratings, slower categorization, and better memory accuracy. Notably, these relationships generally increased in strength from earlier to higher layers. Accordingly, we found no evidence for the reversal reported by Koch *et al.* [26], in which greater distinctiveness in higher layers was associated with reduced memorability.

Together, these findings support the hypothesis that scene typicality – and conversely, distinctiveness – indexes a fundamental property of representational space and that an image’s featural distinctiveness determines both its categorization as a member of a category and its memorability.

### Experiment 3

Experiments 1 and 2 showed that atypical images are remembered more accurately than typical ones, supporting a role of distinctiveness in image memorability. However, it remains unclear which form of distinctiveness drives this advantage: whether atypical images are memorable because they deviate from long-term category schemas, or because they are rare within the specific stimulus set used in the experiment. This distinction has been described as absolute versus relative distinctiveness [51], or as secondary versus primary distinctiveness [13], following William James’ differentiation between long-term and short-term memory.

Accounts that conceptualize memorability as an *intrinsic* image property emphasize its stability across experimental contexts [2, 6], suggesting that memorability reflects an image’s distinctiveness relative to a lifetime of visual experience rather than to the immediate experimental context. Consistent with this view, Bylinskii *et al.* [6] showed that although overall memory performance decreases as the number of category exemplars increases, the rank order of image memorability remains stable. However, most prior studies, including Experiments 1 and 2 of the present work, used stimulus sets with relatively uniform memorability distributions, containing only a small number of highly distinctive images. Under these conditions, memorable images may have benefited from additional relative (primary) distinctiveness by standing out as unusual events within the experiment [10, 12].

To establish that image memorability is truly intrinsic, it is therefore necessary to demonstrate not only robustness to the number of category exemplars [6], but also independence from the relative prevalence of typical and atypical images. Experiment 3 addressed this issue by systematically varying the proportion of atypical images across blocks, allowing us to test whether atypical images retain their memory advantage even when atypicality is no longer rare, i.e. when most of the presented images are atypical.

#### Methods

The methods of the third experiment were the same as for the first two experiments, except for the following changes.

#### Participants

Data were collected from 174 participants (125 female; age range 18-65, mean 24 years; 158 right-handed). Data from 24 participants were excluded because they included less than five correct trials per typicality condition within the range of 200 to 2000 ms, and 33 were excluded because they made false alarms to more than two catch trials, resulting in data from 128 participants used for the analysis.

#### Stimuli and Procedure

Experiment 3 was only conducted online. Each categorization block contained 40 target images from a single scene category. Across three blocks, the proportion of typical images was varied such that each participant completed one block with 75% typical images, one with 50% typical images, and one with 25% typical images. Each categorization block also included 40 distractor images from non-target categories. These distractors were selected so that their typicality ratings were close to the median of their respective categories. The corresponding recognition block included 40 foil images from the same target category, matched to the targets in their proportion of typical and atypical images, and four catch trials from a novel category (stairwell or supermarket). The assignment of scene category and typicality proportion was counterbalanced across participants.

#### Analysis

Data were analyzed using 2 × 3 repeated-measures ANOVAs with the factors *typicality* (high, low) and *proportion* of typical images (rare: 25%, balanced: 50%, frequent: 75%).

#### Results

Categorization and recognition memory performance are summarized in Figure 6.

**Figure 6:**
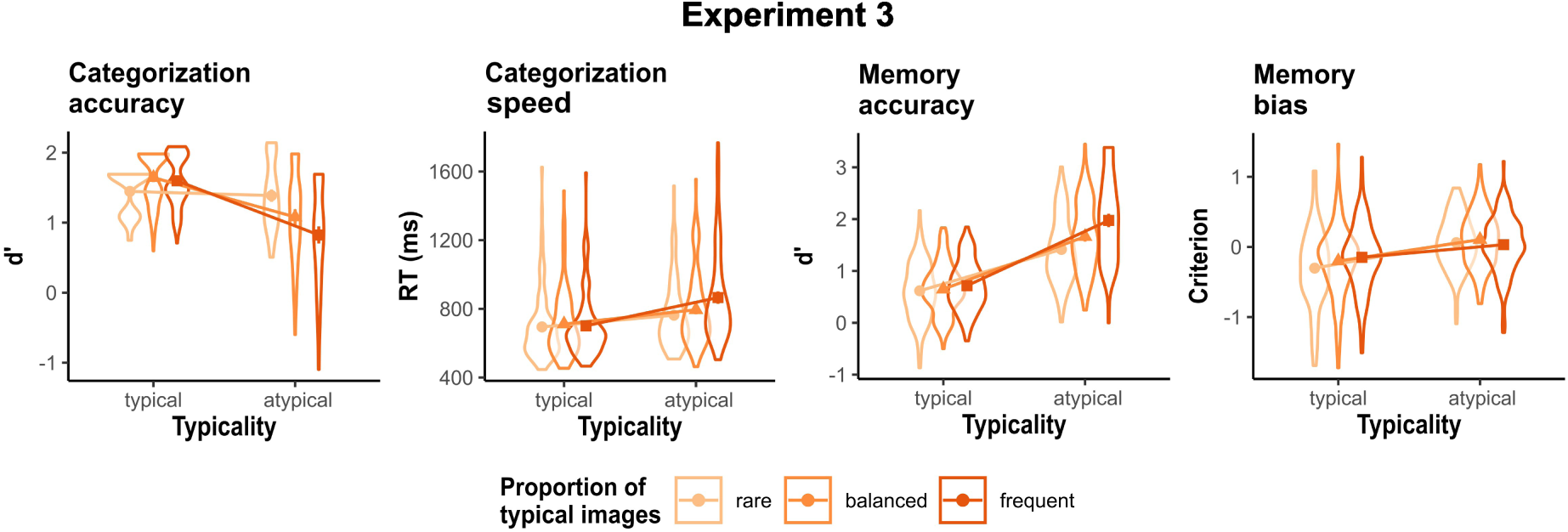
Categorization was more accurate and faster for typical than for atypical scenes, particularly when typical scenes occurred more frequently. By contrast, recognition memory was generally more accurate for atypical scenes. This memory advantage was strongest when the proportion of typical and atypical scenes was balanced and weakest when typical scenes were more frequent. Nevertheless, atypical images were remembered best regardless of their relative proportion. The recognition criterion was lower for typical scenes—especially when typical images were rare, indicating a more liberal bias to judge typical images as “old.”

#### Categorization Task

##### Accuracy

Categorization was more accurate for typical targets than for atypical targets (*F* (1, 127) = 214.08, *p* < .001, 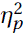 = .63). Accuracy was generally highest when typical targets were frequent, intermediate in the balanced condition, and lowest when typical targets were rare (*F* (2, 254) = 10.09, *p* < .001, 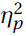 = .07). A significant interaction (*F* (1.85, 234.91) = 36.21, *p* < .001, 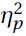 = .22) indicated that the typicality effect on accuracy was stronger when typical targets were frequent (*t* (127) = 11.44, *p <* .001), than in the balanced condition (*t* (127) = 9.43, *p <* .001), and not significant when typical targets were rare (*t* (127) = 1.24, *p* = 0.217).

##### Response times

Categorization was faster for typical than for atypical targets (*F* (1, 127) = 274.39, *p* < .001, 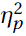 = .68). Response times were fastest when typical targets were frequent, intermediate in the balanced condition, and slowest when typical targets were rare (*F* (1.90, 241.81) = 5.92, *p*=.004, 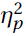 = .04). A significant interaction (*F* (1.54, 196.17) = 18.68, *p* < .001, 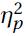 = .13) indicated that the typicality effect on categorization speed was strongest when typical targets were frequent (*t* (127) = −10.03, *p <* .001), intermediate in the balanced condition, (*t* (127) = −9.21, *p <* .001), and weakest when typical targets were rare (*t* (127) = −9.01, *p <* .001).

#### Memory Task

##### Accuracy

Recognition accuracy was higher for atypical images than for typical images (*F* (1, 127) = 681.92, *p* < .001, 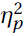 = .84). Moreover, accuracy was highest when typical images were frequent, intermediate for the balanced condition, and lowest when typical images were rare (*F* (2, 254) = 17.87, *p* < .001, 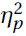 = .12). A significant interaction (*F* (2, 254) = 15.04, *p* < .001, 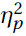 = .11) indicated that the typicality effect on accuracy was strongest when the proportion of typical and atypical images was balanced (*t* (127) = −19.66, *p <* .001), intermediate when typical images were frequent (*t* (127) = −18.39, *p <* .001), and lowest when typical images were rare (*t* (127) = −12.07, *p <* .001).

##### Criterion

Recognition criterion was negative for typical scenes (mean = –0.2174) and positive for atypical scenes (mean = 0.0662), indicating that participants were biased to judge typical scenes as “old”. This liberal bias for typical scenes was confirmed by a main effect of typicality (*F* (1, 127) = 102.96, *p* < .001, 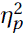 = .45). Moreover, criterion was generally lowest (i.e. most liberal) when typical images were rare, intermediate when typical images were frequent, and lowest in the balanced condition (*F* (2, 254) = 3.68, *p* = .027, 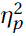 = .03). A significant interaction (*F* (2, 254) = 5.31, *p* = .005, 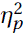 = .04) indicated that the typicality effect on criterion was strongest when typical images were rare, (*t* (127) = −8.37, *p <* .001), intermediate in the balanced condition, (*t* (127) = −7.86, *p <* .001), and lowest when typical images were frequent (*t* (127) = −3.83, *p <* .0002).

## Discussion

Experiment 3 confirmed the main finding of Experiments 1 and 2: atypical images were categorized more slowly but remembered more accurately. By manipulating the relative frequencies of typical and atypical images, we dissociated primary distinctiveness (rarity within the experimental context) from secondary distinctiveness (atypicality relative to long-term category knowledge). If memorability were driven solely by primary distinctiveness, then whichever image type was rarer in a given block should have been remembered best.

Instead, we observed a robust main effect of typicality: memory accuracy was consistently higher for atypical images, regardless of their relative proportion. Although this memory advantage for atypical images was amplified when they were rare – indicating that primary distinctiveness modulates recognition performance – the advantage did not reverse: atypical images were still remembered best even when most images were atypical. Thus, maximizing the primary distinctiveness of typical images did not make them more memorable than atypical images.

In sum, although the results demonstrate that experimental context influences memory performance, they strongly confirm the notion that memorability includes a context-independent *intrinsic* component grounded in secondary distinctiveness.

## General Discussion

A growing body of work has shown that the memorability of scene images is remarkably consistent across observers, suggesting that memorability reflects intrinsic properties of visual stimuli rather than idiosyncratic experiences or task-specific factors. Yet, what makes an image intrinsically memorable remains an open question. Here, we asked whether intrinsic memorability can be understood in terms of distinctiveness: the degree to which an image deviates from a schema or prototype of a learned visual category. A central contribution of the present study is that we operationalize image distinctiveness using three converging measures: subjective typicality judgments, behavioral categorization speed, and computational estimates derived from deep neural network representations. Each of these measures captures a different level at which an image’s relation to its category can be assessed: phenomenological, behavioral, and representational. Focusing on exemplars from a small set of indoor scene categories, we examined how within-category typicality and distinctiveness shape both categorization and recognition memory performance. This approach allowed us to test exemplar-level differences in memorability while minimizing confounds arising from differences in memorability across semantic categories. Note that throughout this paper, “typicality” refers to human judgments about how representative an image is of its semantic category. In contrast, “distinctiveness” refers to a computational property of an image, quantified as the degree to which its visual features are dissimilar from those of other exemplars within the same category, based on CNN-derived feature representations. Building on representational theories of categorization and memory, we tested the hypothesis that an image’s position within a category-specific representational space determines how efficiently it is categorized and how well it is remembered.

Across three experiments, we observed a robust dissociation between categorization performance and recognition memory as a function of scene typicality. Typical scene exemplars were categorized more rapidly and more accurately than atypical exemplars, consistent with the idea that they closely match category-defining features. In contrast, typical scenes were remembered more poorly: they showed lower recognition sensitivity and elicited a more liberal response criterion, indicating an increased tendency to judge new typical scenes as previously seen. Specifically, Experiment 2 demonstrates that the effect of typicality holds not only at the group-level, but also at the level of individual images, where an image’s average categorization speed predicted its memory accuracy.

Critically, Experiment 3 addressed a central challenge to interpreting memorability as an intrinsic image property: the possibility that atypical images benefit simply from standing out within the experimental context. Previous studies have varied the number of exemplars per category and found that overall memory performance decreases slightly, while the rank order of image memorability remains stable – an observation often taken as evidence that memorability is stable across experimental contexts [6, 27, 52]. However, to our knowledge, this is the first study directly contrasting image typicality with primary distinctiveness by manipulating the proportions of memorable versus non-memorable or distinctive versus non-distinctive images within an experiment. This distinction is critical because some situational factors, such as the total number of exemplars or presentation order, may only influence overall memory performance while remaining orthogonal to intrinsic memorability. In contrast, the distribution of memorability *within* the stimulus set could fundamentally confound estimates of intrinsic memorability. Specifically, many memorability experiments rely on convenience samples of images yielding an approximately uniform or normal distribution of memorability, making highly memorable images relatively rare and stand out strongly from the experimental context. These images may be remembered better not because they are intrinsically distinctive relative to a lifetime of visual experience (secondary distinctiveness), but because they are distinctive relative to the other images shown in the experiment (primary distinctiveness, [13]). To address this issue, Experiment 3 systematically manipulated the frequency of atypical, and thus distinctive images. While recognition accuracy generally improved when atypical images were rare, consistent with an effect of primary distinctiveness, we observed that a robust recognition advantage for atypical images persists even when most images are atypical. To our knowledge, this is the first study to directly contrast image typicality with primary distinctiveness. This finding indicates that the memory advantage for atypical scenes cannot be explained solely by their primary distinctiveness within the short-term experimental context, but instead reflects representational distinctiveness arising from their distal position in category-specific representational space, based on long-term visual experience.

Taken together, these results support the view that scene memorability is governed by representational distinctiveness: the degree to which an image occupies a peripheral versus central position within a learned category-specific feature space. This idea is not new in principle. Early theories of distinctiveness conceptualized items as points in high-dimensional feature spaces, with typical category members forming dense clusters near the center and atypical exemplars residing in sparser, more isolated regions [15, 53]. What has historically been lacking, however, is a principled way to quantify such representational distances for complex natural scenes. Here, we address this gap by linking three related indicators of distinctiveness: subjective typicality judgments, categorization speed, and computational estimates derived from deep neural network representations. Importantly, the neural network we used was trained solely to perform scene categorization, not to predict memorability, typicality, or response times. Its ability to account for these behavioral measures, and therefore, does not reflect task-specific optimization, but instead arises because it captures feature representations that approximate the structure of human visual categories. In this sense, CNN-derived distinctiveness serves as a quantitative proxy for the geometry of representational space, supporting the view that memorability reflects an image’s relational position within a category-specific representational space.

Analyses of layer-specific representations in the deep neural network further constrain how distinctiveness relates to memorability. Distinctiveness derived from early convolutional layers showed only weak relationships with typicality, categorization speed, and recognition memory. This pattern is consistent with prior work showing that memorability is only weakly related to low-level visual properties such as luminance, contrast, or color [1, 4]. In contrast, stronger and more systematic relationships emerged at intermediate and higher layers of the network. Categorization speed showed its strongest relationship with distinctiveness derived from an intermediate convolutional layer, suggesting that the efficiency of category verification is most tightly linked to mid-level visual representations that capture diagnostic feature combinations. In contrast, recognition memory and subjective typicality were most strongly correlated with distinctiveness in later, more category-selective layers of the network. The present results therefore suggest that memorability is linked not to low-level features or how unusual an image looks, but to how an image is embedded within higher-level, category-relevant representations.

Previous work has addressed the relationship between distinctiveness and memory, but the empirical picture has remained mixed [54]. In line with our findings, research on face memory has consistently shown that atypical faces are remembered better than typical faces, based both on subjective typicality ratings and on computational measures of feature distinctiveness. Within norm-based face space models, this advantage is explained by the greater representational distance of distinctive faces from the average face or prototype [55]. Importantly, mirroring our findings for scenes, atypical faces – despite being easier to remember – are harder to categorize than typical faces [24].

A recent study on scene memory examined scene memorability across a broad range of semantic categories using a computational measure of feature distinctiveness derived from a deep neural network [26]. Scenes whose features were distinctive in the first convolutional layer were more memorable, whereas scenes that were distinctive in the final classification layer (fc8) were less memorable. At first glance, this pattern appears to conflict with our results, which show greater memorability for distinctive images across all layers, with the weakest association observed in the earliest convolutional layer (Figures 4, 5). A key difference between the two studies lies in the stimulus set and how distinctiveness was defined. In the present work, distinctiveness was computed only among exemplars from the same semantic category, such that images with low distinctiveness closely resembled the category prototype. From this perspective, it is not surprising that prototypical images – whose features are similar to many other exemplars – were more confusable in memory. By contrast, Koch *et al.* [26] computed distinctiveness across images drawn from many different scene categories, with only a small number of exemplars per category. Under these conditions, high similarity to other images does not clearly map onto category typicality, making it more difficult to interpret what “distinctive” or “non-distinctive” images represent at higher, category-selective layers.

Kramer *et al.* [27] investigated the memorability of object images from the THINGS database, which comprises images of objects associated with semantic categories. Object features derived from human similarity judgments showed only a weak relationship with memorability, such that more distinctive images tended to be less memorable. Similarly, images with more distinctive features derived from either early or higher CNN layers were, on average, less memorable, although this relationship was highly heterogeneous across object categories. Surprisingly, human typicality judgments were not correlated with memorability at all. In contrast, our results show the opposite pattern: distinctiveness in both early and late network layers strongly predicts higher memorability, and scenes judged as atypical are likewise remembered better.

One potentially relevant difference between the studies concerns how typicality was operationalized. In Kramer *et al.* [27], typicality ratings reflect how well the depicted concept fits a superordinate category (e.g., whether a snake belongs to the category “animal”). By contrast, our images were rated for typicality at the basic level – for example, not whether a kitchen is a typical room, but whether a given kitchen image shows a typical kitchen. The studies also differed in their stimuli: most images in the THINGS database depict a single foreground object that can be easily named (e.g., “snake”) and are drawn from a broad range of semantic concepts (e.g., animal), whereas our stimuli consisted of complex scenes containing multiple objects.

Recently, several authors have proposed that memorable images form stronger memories because they are processed more efficiently at early perceptual stages [54, 56]. Consistent with this view, memorable images can be identified with shorter presentation durations [57] and show greater resistance to visual masking [5, 56]. These findings have motivated the proposal that the enhanced perceptual processing of memorable images arises because they are statistically more regular [56] or more closely match a category template or prototype than less memorable images [54]. If this account is correct, memorable images should therefore be more typical and easier to categorize. Our results do not support this prediction. In our data, memorable images showed a disadvantage in categorization performance and were overall less typical than non-memorable images. Together, these findings suggest that typicality and perceptual processing efficiency may constitute partially independent determinants of image memorability.

A key design choice of the present study was to focus on only three indoor scene categories. This restricted stimulus set was intentional and constitutes a strength of the approach, as it allowed us to isolate exemplar-level typicality effects while minimizing confounds arising from general category differences in semantics or baseline memorability. Notably, the effects of typicality were highly consistent across the three categories, suggesting that the observed relationships were not driven by idiosyncratic properties of a single category. At the same time, this choice raises questions about generalizability. It remains unclear whether a similar memorability benefit for atypical exemplars would emerge in paradigms that include many categories but only a few exemplars per category – a design common in large-scale memorability studies. One reason is that distinctiveness is inherently context-dependent: an item is distinctive only relative to a set of similar items [10, 53]. In our design, the presence of many exemplars from the same category created a meaningful reference set in which atypical scenes could stand out from more typical ones. By contrast, when stimuli are drawn from many different categories with few exemplars per category, this within-category reference structure may be weaker, potentially reducing the influence of exemplar-level distinctiveness.

Moreover, our stimuli were limited to man-made indoor environments, which are shaped by explicit functional and cultural category structures. For example, architectural conventions and regulations impose clear constraints on what constitutes a “kitchen” or “bathroom”, potentially reinforcing strong category prototypes. Natural environments, by contrast, are not organized around such normative definitions, and category boundaries for natural scenes such as mountains, forests, or lakes may be more graded or ambiguous. Whether exemplar typicality predicts memorability in a comparable manner for natural scene categories therefore remains an open empirical question. Addressing these issues in future work will be essential for determining how the relationship between image typicality and memorability generalizes across stimulus domains and experimental contexts.

In sum, our findings support a representational account of image memorability in which memory is determined by an image’s position within a learned, category-specific representational space. By integrating subjective typicality judgments, categorization performance, and deep neural network representations, we show that atypical scene exemplars are remembered better because they are representationally distinctive relative to category prototypes shaped by long-term visual experience.

## Acknowledgements

This work was supported by Deutsche Forschungsgemeinschaft (DFG) Grant BU2400/14-1 and by Czech Science Foundation Grant 24-11506K.

## References

1. Isola, P., Xiao, J., Torralba, A. & Oliva, A. What makes an image memorable? in CVPR 2011 (2011), 145–152. doi:10.1109/CVPR.2011.5995721.

2. Bainbridge, W. A. Memorability: Reconceptualizing memory as a visual attribute in Visual Memory (Routledge, 2022).

3. Bylinskii, Z., Goetschalckx, L., Newman, A. & Oliva, A. Memorability: An Image-Computable Measure of Information Utility in Human Perception of Visual Information: Psychological and Computational Perspectives 207–239 (Springer International Publishing, Cham, 2022). doi:10.1007/978-3-030-81465-6_8.

4. Isola, P., Xiao, J., Parikh, D., Torralba, A. & Oliva, A. What Makes a Photograph Memorable? IEEE Transactions on Pattern Analysis and Machine Intelligence 36, 1469–1482. doi:10.1109/TPAMI.2013.200 (2014).

5. Goetschalckx, L., Moors, P., Vanmarcke, S. & Wagemans, J. Get the Picture? Goodness of Image Organization Contributes to Image Memorability. Journal of Cognition 2, 22. doi:10.5334/joc.80 (2019).

6. Bylinskii, Z., Isola, P., Bainbridge, C., Torralba, A. & Oliva, A. Intrinsic and extrinsic effects on image memorability. Vision Research 116, 165–178. doi:10.1016/j.visres.2015.03.005 (2015).

7. Goetschalckx, L. & Wagemans, J. MemCat: a new category-based image set quantified on memorability. PeerJ 7, e8169. doi:10.7717/peerj.8169 (2019).

8. Khosla, A., Raju, A. S., Torralba, A. & Oliva, A. Understanding and Predicting Image Memorability at a Large Scale in 2015 IEEE International Conference on Computer Vision (ICCV) (IEEE, Santiago, Chile, 2015), 2390–2398. doi:10.1109/ICCV.2015.275.

9. Needell, C. D. & Bainbridge, W. A. Embracing New Techniques in Deep Learning for Estimating Image Memorability. Computational Brain & Behavior 5, 168–184. doi:10.1007/s42113-022-00126-5 (2022).

10. Hunt, R. R. The subtlety of distinctiveness: What von Restorff really did. Psychonomic Bulletin & Review 2, 105–112. doi:10.3758/BF03214414 (1995).

11. Konkle, T., Brady, T. F., Alvarez, G. A. & Oliva, A. Conceptual distinctiveness supports detailed visual long-term memory for real-world objects. Journal of Experimental Psychology. General 139, 558–578. doi:10.1037/a0019165 (2010).

12. Von Restorff, H. Über die Wirkung von Bereichsbildungen im Spurenfeld. Psychologische Forschung 18, 299–342. doi:10.1007/BF02409636 (1933).

13. Schmidt, S. R. Can we have a distinctive theory of memory? Memory & Cognition 19, 523–542. doi: 10.3758/BF03197149 (1991).

14. Rosch, E., Simpson, C. & Miller, R. S. Structural bases of typicality effects. Journal of Experimental Psychology: Human Perception and Performance 2, 491–502. doi:10.1037/0096-1523.2.4.491 (1976).

15. Schmidt, S. R. Category typicality effects in episodic memory: Testing models of distinctiveness. Memory & Cognition 24, 595–607. doi:10.3758/BF03201086 (1996).

16. Friedman, A. Framing pictures: The role of knowledge in automatized encoding and memory for gist. Journal of Experimental Psychology: General 108, 316–355. doi:10.1037/0096-3445.108.3.316 (1979).

17. Going, M. & Read, J. D. Effects of Uniqueness, Sex of Subject, and Sex of Photograph on Facial Recognition. Perceptual and Motor Skills 39, 109–110. doi:10.2466/pms.1974.39.1.109 (1974).

18. Light, L. L., Kayra-Stuart, F. & Hollander, S. Recognition memory for typical and unusual faces: Journal of Experimental Psychology: Human Learning and Memory. Journal of Experimental Psychology: Human Learning and Memory 5, 212–228. doi:10.1037/0278-7393.5.3.212 (1979).

19. Vokey, J. R. & Read, J. D. Familiarity, memorability, and the effect of typicality on the recognition of faces. Memory & Cognition 20, 291–302. doi:10.3758/bf03199666 (1992).

20. Rakover, S. S. & Cahlon, B. Face recognition: cognitive and computational processes Advances in consciousness research 31 (John Benjamins Pub. Co, Amsterdam Philadelphia, PA, 2001).

21. Lukavský, J. & Děchtěrenko, F. Visual properties and memorising scenes: Effects of image-space sparseness and uniformity. *Attention, Perception*, & Psychophysics 79, 2044–2054. doi:10.3758/s13414-017-1375-9 (2017).

22. Reed, S. K. Pattern recognition and categorization. Cognitive Psychology 3, 382–407. doi:10.1016/0010-0285(72)90014-X (1972).

23. Smith, E. E., Shoben, E. J. & Rips, L. J. Structure and process in semantic memory: A featural model for semantic decisions. Psychological Review 81, 214–241. doi:10.1037/h0036351 (1974).

24. Valentine, T. & Bruce, V. The Effects of Distinctiveness in Recognising and Classifying Faces. Perception 15, 525–535. doi:10.1068/p150525 (1986).

25. Torralbo, A. et al. Good Exemplars of Natural Scene Categories Elicit Clearer Patterns than Bad Exemplars but Not Greater BOLD Activity. PLoS ONE 8, e58594. doi:10.1371/journal.pone.0058594 (2013).

26. Koch, G. E., Akpan, E. & Coutanche, M. N. Image memorability is predicted by discriminability and similarity in different stages of a convolutional neural network. Learning & Memory (Cold Spring Harbor, N.Y.) 27, 503–509. doi:10.1101/lm.051649.120 (2020).

27. Kramer, M. A., Hebart, M. N., Baker, C. I. & Bainbridge, W. A. The features underlying the memorability of objects. Science Advances 9, eadd2981. doi:10.1126/sciadv.add2981 (2023).

28. Peirce, J. et al. PsychoPy2: Experiments in behavior made easy. Behavior Research Methods 51, 195–203. doi:10.3758/s13428-018-01193-y (2019).

29. Sareen, P., Ehinger, K. A. & Wolfe, J. M. CB Database: A change blindness database for objects in natural indoor scenes. Behavior Research Methods 48, 1343–1348. doi:10.3758/s13428-015-0640-x (2016).

30. Xiao, J., Hays, J., Ehinger, K. A., Oliva, A. & Torralba, A. SUN database: Large-scale scene recognition from abbey to zoo in 2010 IEEE Computer Society Conference on Computer Vision and Pattern Recognition (2010), 3485–3492. doi:10.1109/CVPR.2010.5539970.

31. Green, D. M. & Swets, J. A. Signal Detection Theory and Psychophysics (Wiley, 1966).

32. Hautus, M. J. Corrections for extreme proportions and their biasing effects on estimated values of d. Behavior Research Methods, Instruments, & Computers 27, 46–51. doi:10.3758/BF03203619 (1995).

33. Singmann, H., Bolker, B., Westfall, J., Aust, F. & Ben-Shachar, M. S. afex: Analysis of factorial experiments manual (2021).

34. R Core Team. R: A language and environment for statistical computing manual (Vienna, Austria, 2022).

35. Lenth, R. V. emmeans: Estimated marginal means, aka least-squares means manual (2021).

36. Rosch, E. H. On the internal structure of perceptual and semantic categories in Cognitive Development and Acquisition of Language 111–144 (Academic Press, San Diego, 1973). doi:10.1016/B978-0-12-505850-6.50010-4.

37. Rips, L. J., Shoben, E. J. & Smith, E. E. Semantic distance and the verification of semantic relations. Journal of Verbal Learning and Verbal Behavior 12, 1–20. doi:10.1016/S0022-5371(73)80056-8 (1973).

38. Nosofsky, R. M. Attention, similarity, and the identification-categorization relationship. Journal of Experimental Psychology. General 115, 39–61. doi:10.1037//0096-3445.115.1.39 (1986).

39. Delhaye, E., Coco, M. I., Bahri, M. A. & Raposo, A. Typicality in the brain during semantic and episodic memory decisions. Neuropsychologia 184, 108529. doi:10.1016/j.neuropsychologia.2023. 108529(2023).

40. Krizhevsky, A., Sutskever, I. & Hinton, G. E. ImageNet Classification with Deep Convolutional Neural Networks in Advances in Neural Information Processing Systems 25 (Curran Associates, Inc., 2012).

41. Zeiler, M. D. & Fergus, R. Visualizing and Understanding Convolutional Networks in Computer Vision – ECCV 2014 (Springer International Publishing, Cham, 2014), 818–833. doi:10.1007/978-3-319-10590-1_53.

42. Yamins, D. L. K. & DiCarlo, J. J. Using goal-driven deep learning models to understand sensory cortex. Nature Neuroscience 19, 356–365. doi:10.1038/nn.4244 (2016).

43. Yamins, D. L. K. et al. Performance-optimized hierarchical models predict neural responses in higher visual cortex. Proceedings of the National Academy of Sciences 111, 8619–8624. doi:10.1073/pnas.1403112111 (2014).

44. Eickenberg, M., Gramfort, A., Varoquaux, G. & Thirion, B. Seeing it all: Convolutional network layers map the function of the human visual system. NeuroImage 152, 184–194. doi:10.1016/j.neuroimage.2016.10.001 (2017).

45. Lindsay, G. W. Convolutional Neural Networks as a Model of the Visual System: Past, Present, and Future. Journal of Cognitive Neuroscience 33, 2017–2031. doi:10.1162/jocn_a_01544 (2021).

46. Peterson, J. C., Abbott, J. T. & Griffiths, T. L. Evaluating (and Improving) the Correspondence Between Deep Neural Networks and Human Representations. Cognitive Science 42, 2648–2669. doi:10.1111/cogs.12670 (2018).

47. Brady, T. F. & Störmer, V. S. Comparing memory capacity across stimuli requires maximally dissimilar foils: Using deep convolutional neural networks to understand visual working memory capacity for real-world objects. Memory & Cognition 52, 595–609. doi:10.3758/s13421-023-01485-5 (2024).

48. Paszke, A. et al. PyTorch: An Imperative Style, High-Performance Deep Learning Library 2019. doi:10.48550/arXiv.1912.01703.

49. Zhou, B., Lapedriza, A., Khosla, A., Oliva, A. & Torralba, A. Places: A 10 Million Image Database for Scene Recognition. IEEE Transactions on Pattern Analysis and Machine Intelligence 40, 1452–1464. doi:10.1109/TPAMI.2017.2723009 (2018).

50. Lake, B. M., Zaremba, W., Fergus, R. & Gureckis, T. M. Deep Neural Networks Predict Category Typicality Ratings for Images. Proceedings of the Annual Meeting of the Cognitive Science Society 37 (2015).

51. Hosie, J. A. & Milne, A. B. Distinctiveness and memory for unfamiliar faces in Cognitive and Computational Aspects of Face Recognition (Routledge, 1995).

52. Konkle, T., Brady, T. F., Alvarez, G. A. & Oliva, A. Scene memory is more detailed than you think: the role of categories in visual long-term memory. Psychological Science 21, 1551–1556. doi:10.1177/0956797610385359 (2010).

53. Murdock, B. B. J. The distinctiveness of stimuli. Psychological Review 67, 16–31. doi:10.1037/h0042382 (1960).

54. Bainbridge, W. A., Walther, D. B., Fukuda, K. & Goetschalckx, L. Memorability of visual stimuli and the role of processing efficiency. Nature Reviews Psychology 5, 47–58. doi:10.1038/s44159-025-00512-3 (2026).

55. Valentine, T., Lewis, M. B. & Hills, P. J. Face-Space: A Unifying Concept in Face Recognition Research. Quarterly Journal of Experimental Psychology 69, 1996–2019. doi:10.1080/17470218.2014.990392 (2016).

56. Deng, W., Federmeier, K. D. & Beck, D. M. Highly memorable images are more readily perceived. Journal of Experimental Psychology. General 153, 1415–1424. doi:10.1037/xge0001594 (2024).

57. Broers, N., Potter, M. C. & Nieuwenstein, M. R. Enhanced recognition of memorable pictures in ultra-fast RSVP. Psychonomic Bulletin & Review 25, 1080–1086. doi:10.3758/s13423-017-1295-7 (2018).

